# Positional and Compositional Analysis of Saturated, Monounsaturated and Polyunsaturated Fatty acids in Human Adipose Tissue Triglyceride by ^13^C NMR

**DOI:** 10.1101/2021.02.25.432655

**Authors:** Alejandra N. Torres, Ludgero Tavares, Maria J. Pereira, Jan W. Eriksson, John G. Jones

**Affiliations:** Metabolism, Aging and Disease, Center for Neurosciences and Cell Biology, University of Coimbra, Cantanhede, Portugal; Department of Medical Sciences, Clinical Diabetes and Metabolism, Uppsala University, Uppsala, Sweden

**Keywords:** Triglycerides, fatty acids, palmitoleic acid, ^13^C nuclear magnetic resonance, adipose tissue, metabolism, distribution, BMI, HOMA

## Abstract

**Objectives and new findings, ends with short conclusion, no references:** The synthesis and turnover of triglyceride in adipose tissue involves enzymes with preferences for specific fatty acid classes and/or regioselectivity with regard to the fatty acid position within the glycerol moiety. The focus of the present study was to characterize both the fatty acid composition and their positional distribution in triglycerides of biopsied human subcutaneous adipose tissue using ^13^C NMR spectroscopy. The triglyceride *sn*2 position was significantly more enriched with monounsaturated fatty acids compared to the *sn*1,3 sites, while saturated fatty acids abundance was significantly lower in the *sn*2 position compared to that of *sn*1,3. Furthermore, the analysis revealed significant positive correlations between the total fraction of palmitoleic acid with both BMI and HOMA-IR scores. Additionally, we established that ^13^C NMR chemical shifts for ω −3 signals, centered at 31.9 ppm, provided superior resolution of the most abundant FA species, including palmitoleate, compared to the ω −2 signals that were used previously. ^13^C NMR spectroscopy reveals for the first time a highly non-homogenous distribution of FA in the glycerol sites of human adipose tissue triglyceride and that these distributions are correlated with different phenotypes such as BMI and insulin resistance.

## Introduction

The canonical function of adipose tissue triglycerides (TG) is to provide essential and non-essential fatty acids (FA) to other tissues as energy substrates and as precursors for the biosynthesis of cellular components such as phospholipids (1). Certain FA species such (16:1, *n-*7) may also contribute to metabolic regulation and insulin sensitivity in other tissues thereby involving the adipose tissue in the coordination of metabolic homeostasis for the whole body (2). In the setting of obesity and insulin resistance, the overall FA composition of adipose tissue TG may be influenced by changes in *de novo* FA synthesis, elongation and desaturation as well as by dietary lipid constitution (3,4,5,6).

TG synthesis involves the sequential addition of fatty acids to the *sn-1, sn-2* and *sn-3* positions of glycerol (7) by glycerol-3-P acyltransferase, monoacylglycerol-3-P acyltransferase and diacylglycerol acyltransferase - each of which has a different affinity for saturated fatty acids (SFA), monounsaturated fatty acids (MUFA) and polyunsaturated fatty acids (PUFA) (8,9,10). Likewise, TG hydrolysis is also a coordinated process involving adipose triglyceride lipase, hormone-sensitive lipase and monoacylglycerol lipase - each of which also has selective affinity for different fatty acid classes (11,12). While net TG synthesis and lipolysis are reciprocally regulated according to fed and fasted states, the two processes can occur simultaneously, resulting in cycling between TG and FA (13,14). To the extent that these activities are modified by disease states, the positional distribution of different FA species in the TG molecule may be altered concurrently or independently of FA composition.

Of the methodologies that are in current use for characterizing TG composition, ^13^C NMR is unique in providing information on the positional distribution of FA between the glycerol *sn*1,3 and *sn*2 sites as well as their overall composition (15). We previously characterized FA composition and positional distribution in TG of adipose tissue from healthy mice by ^13^C NMR (16) and found that different FA classes (SFA, MUFA, PUFA) were not uniformly distributed between the glycerol *sn*1,3 and *sn*2 sites. To our knowledge, such observations have not been made for human adipose tissue (AT). Therefore, the aim of this study was to characterize FA composition and positional distribution in TG of biopsied human adipose tissue. Moreover, we studied a subject cohort with wide ranges of body mass index (BMI) and insulin sensitivity to determine if FA composition and/or distribution were correlated with these parameters.

## Methods

### Materials

Synthetic triglycerides of linoleate, linolenate, myristate, oleate, palmitate, palmitoleate, and stearate, chloroform containing amylenes as preservative for triglyceride extraction and deuterated chloroform for triglyceride ^1^H and ^13^C NMR analysis were all obtained from Merck (Merck Portugal, Algés, Portugal).

### Adipose tissue biopsies and Ethics statement

Subcutaneous adipose tissue needle biopsies were obtained from the lower abdomen after local anesthesia with lidocaine (Xylocain, Astrazeneca, Sweden). Following adipose tissue biopsy, the samples were immediately snap frozen in liquid nitrogen and stored at −80°C. 39 healthy controls and 7 subjects with type 2 diabetes were recruited at the Uppsala University Hospital. All participants were >18 years, and subjects with type 1 diabetes, endocrine disorders, cancer or other major illnesses were excluded. The T2D subjects were on a stable dose of metformin for at least the past three months as their only anti-diabetic medication and subjects taking systemic glucocorticoids, beta blockers, and immune modulating therapies, were excluded from the study. Fasting blood samples from study participants were obtained for biochemical profiling (Table 1) at the Department of Clinical Chemistry, Uppsala University Hospital. The study was approved by the Regional Ethics Review Boards in Uppsala and all participants gave their written informed consent. The mass of tissue biopsies that were analyzed by NMR varied from 11.6-65.3 mg.

**Table 1.** Characteristics of the subjects involved in the study. Data are shown as mean ± SD and include Homeostatic model assessment of insulin resistance index (HOMA IR).

| *Characteristics* | *ALL*<br>*Participants* | *Female* | *Male* |
| --- | --- | --- | --- |
| *number* | 46 | 26 | 20 |
| *Age (years)* | 48 ± 17 | 49 ± 15 | 47 ± 20 |
| *Clinically confirmed T2D (n)* | 7 | 3 | 4 |
| *Body mass index (kg/m<sup>2</sup>)* | 30.20 ± 4.87 | 30.90 ± 5.13 | 29.30 ± 4.48 |
| *HOMA IR (mmol x mU/L<sup>2</sup>)* | 3.25 ± 2.10 | 2.88 ± 1.82 | 3.72 ± 2.40 |
| *Plasma Glucose (mmol/L)* | 6.00 ± 1.17 | 5.65 ± 0.78 | 6.46 ± 1.44 |
| *Plasma Cholesterol (mmol/L)* | 5.13 ± 1.17 | 5.17 ± 1.02 | 5.09 ± 1.37 |
| *Plasma HDL-Cholesterol (mmol/L)* | 1.32 ± 0.38 | 1.50 ± 0.40 | 1.10 ± 0.19 |
| *Plasma LDL-Cholesterol (mmol/L)* | 3.32 ± 0.90 | 3.29 ± 0.83 | 3.36 ± 1.00 |
| *Plasma (LDL/HDL)* | 2.60 ± 0.87 | 2.23 ± 0.67 | 3.07 ± 0.89 |
| *Plasma Triglycerides (mmol/L)* | 1.17 ± 0.59 | 1.03 ± 0.45 | 1.36 ± 0.70 |
| *Serum Insulin (mE/L)* | 11.77 ± 6.50 | 11.22 ± 6.31 | 12.49 ± 6.84 |

### Biopsy TG extraction and purification

Triacylglycerols were extracted with methyl-*tert*-butyl ester (MTBE) (17). Methanol (4.6 mL/g) was added to frozen samples and vortexed followed by MTBE (15.4 mL/g) and allowed to stand for 1h at room temperature. The mixture was then centrifuged for 10 min at 13,000 x *g*. The supernatant was collected and phase separation was induced by addition of distilled water (4mL/g) and letting the samples rest for 10 min at room temperature. Phase separation was facilitated by centrifuging samples for 10 min at 1000 x *g*, after which the upper layer containing the lipids was transferred to vials and dried under N_2_ gas.

### NMR Spectroscopy

Purified triacylglycerol samples were dissolved in 0.2 mL 99.96% enriched CDCl_3_ (Sigma-Aldrich). All ^13^C NMR spectra were obtained with a Varian VNMRS 600 MHz NMR (Agilent) spectrometer equipped with a 3 mm standard probe. ^13^C NMR spectra were acquired at 25 °C using a 60° pulse, 30.5 kHz spectral width and 4.1 s of recycling time (4.0 s of acquisition time and 0.1 s pulse delay). The number of free-induction decays (f.i.d.) ranged from 10,000 to 20,000. The spectra were processed with 0.2 Hz line-broadening before Fourier transformation. ^1^H NMR spectra of the same samples were acquired with the same probe and conditions using a 45° pulse, 7 KHz spectral width, and 5.0 s of recycling time (4 s of acquisition time and 1s pulse delay) with 16 f.i.d. per sample. F.i.d. were processed with 0.2 Hz of line broadening and zero-filling to 64 K points. The relative areas of signals in the ^13^C NMR spectra were analyzed using the curve-fitting routine supplied with the ACDLabs 1D NMR processor software. Relative areas of fatty ^1^H NMR signal clusters were quantified by manual integration using the NUTS software (NUTS, Acorn labs).

### Quantification of triglyceride fatty acid components

The ^13^C chemical shifts of FA carbons are sensitive to their overall concentration in the NMR sample (18). Therefore, ^13^C-chemical shifts of individual FAs and/or FA classes were measured with a standard sample consisting of a mixture of seven TG based on the reported composition of TG FAs in human adipose tissues (4) and their commercial availability. The FA composition of the standard was myristic (14:0, 2.35 mol%), palmitic (16:0, 2.62 mol%), palmitoleic (16:1 n-7, 5.98 mol%), stearic (18:0, 3.72 mol%), oleic (18:1, n-9, 44.91 mol%), linoleic (18:2 n-6, 12.99 mol%) and α-linolenic (18:3 n-3, 0.63 mol%). To quantify the fraction of SFA (defined as the sum of 14:0, 16:0 and 18:0 fatty acids), oleate (OL), palmitoleate (PO) and linoleate (LO), their ω-3 ^13^C signals clustered at 31-32 ppm were measured. From the ^13^C carboxyl signals, the relative fractions of SFA, MUFA and PUFA were measured in the *sn*1,3 and *sn*2 sites (15,16). For both carboxyl and aliphatic ^13^C signals, no corrections were made for differential longitudinal relaxation times or nuclear Overhauser effects. The fraction of ω-3 polyunsaturated fatty acids (ω-3-PUFA), whose ω-3 ^13^C NMR signals are dispersed downfield amidst those of other olefinic carbons, and in any case have insufficient signal-to-noise ratios for reliable quantification, were measured from the ^1^H NMR spectrum. The intensity of the ω-3-PUFA methyl triplet, centered at 0.98 ppm was measured relative to that of all other fatty acid methyls, centered at 0.85 ppm. The integral for the ω-3-PUFA methyl triplet was corrected for the presence of the upfield component of the natural abundance ^13^C-satellite signals from the other fatty acids methyls by subtraction of the value corresponding to 0.0055 × integral of ^12^C other fatty acid methyls. The ^13^C NMR signal areas representing saturated fatty acids, oleate, palmitoleate and linoleate were adjusted for the presence of ω-3-PUFA by multiplication with 1-*f*_ω-3-PUFA_, where *f*_ω-3-PUFA_ is the fraction of ω-3-PUFA estimated from the ^1^H NMR spectrum.

### Data presentation and statistical analysis

Data from different subject categories (BMI, HOMA-IR quartiles) are represented as box and whisker plots with the box spanning the 25^th^ to 75^th^ percentiles the whiskers extending to the lowest and highest value and the center line representing the median value. Statistical significance between data sets was tested using simple linear regressions and correlations were tested using Pearson correlation coefficients, one-way ANOVA or two-way ANOVA where appropriate followed by Tukey’s multiple comparisons with the threshold defined by a *p* value < 0.05, using Prism 9 software.

## Results

### Spectra characteristics

Chloroform extracts of the AT biopsies yielded ^13^C and ^1^H NMR spectra with good resolution of TG fatty acid hydrogens and carbons, as shown by Figures 1 and 2. Examination of the fatty acid ^13^C NMR signals from both human AT biopsies as well the TG standard mixture revealed that the ω −3 signals, whose chemical shifts are centered at 31.9 ppm, provided superior resolution of the most abundant FA species compared to the ω −2 signals that were used previously (16) as seen in Figure 1a and Supplementary Figure 1a. ^13^C NMR spectra of individual glyceryl trimyristate, glyceryl tripalmitate and glyceryl tristearate standards showed identical chemical shift values for the ω −3 position (Supplementary Table 1). In the TG standard mixture, they contributed to a single sharp signal representing their summed contribution to the TG fatty acid fraction (Supplementary Figure 1). The fraction of ω −3 polyunsaturated fatty acids (ω −3 PUFA) determined from analysis of the ^1^H NMR spectra accounted for about 2% of the total. For the ^1^H NMR spectrum, the upfield ^13^C-natural abundance satellites of the much larger neighboring methyl signal representing the remaining 98% of triglyceride fatty acids was a significant contributor (20-30%) to the integral used to quantify ω −3 PUFA (Figure 2, Supplementary Figure 2). This was accordingly corrected for, as described in the Methods section.

**Figure 1a,b.**
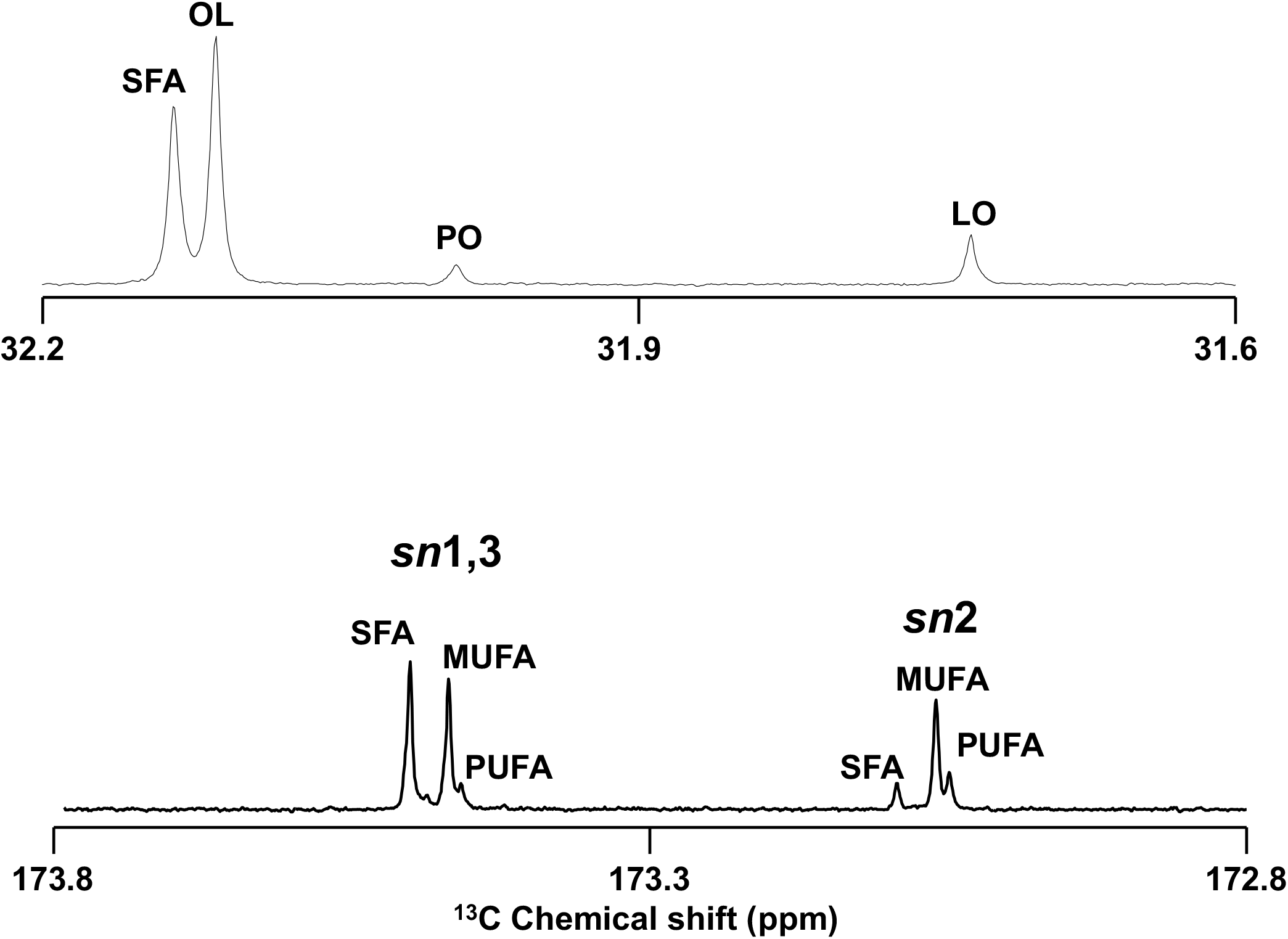
Expanded portions of a ^13^C NMR spectrum obtained from a 21.7 mg adipose tissue biopsy sample. Signals of the ω-3 carbons for saturated fatty acids (SFA), oleic acid (OL), palmitoleic acid (PO) and linoleic acid (LO) are shown in **(a)** while those representing SFA, monounsaturated fatty acids (MUFA) and polyunsaturated fatty acids (PUFA) distributed among the *sn*1,3 and *sn*2 carboxyls are shown in **(b)**.

**Figure 2.**
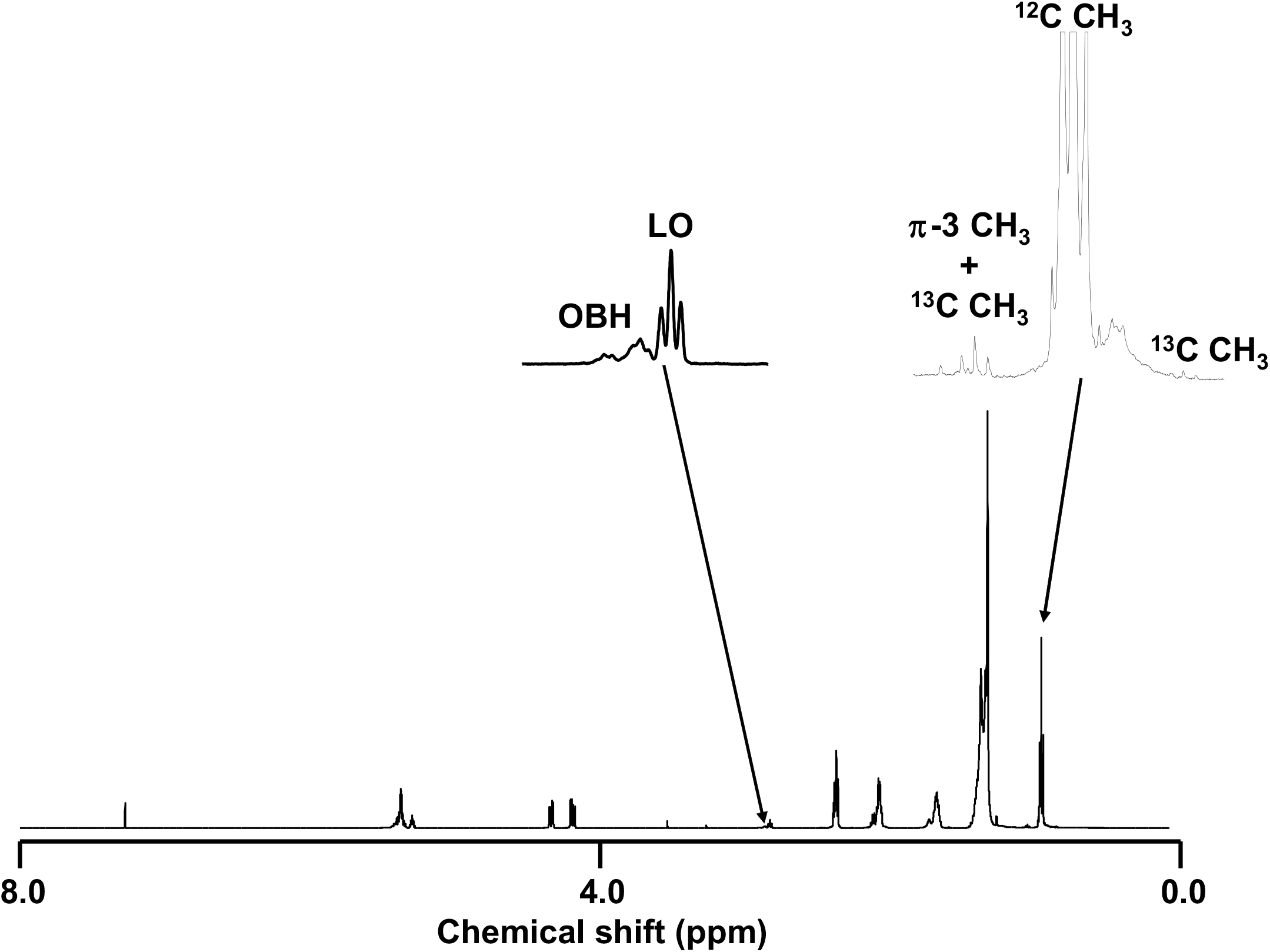
The corresponding ^1^H NMR spectrum of the sample with the fatty acid methyl signals of ω-3 polyunsaturated fatty acids (ω-3 CH_3_) and those of all other fatty acids (^12^C CH_3_) including their ^1^H-^13^C-coupled satellites from ^13^C natural abundance (^13^C CH_3_). Also highlighted are the linoleic acid bisallylic hydrogens (LO). The poorly defined multiplet component upfield of LO represents other bisallylic hydrogens (OBH) from other species of polyunsaturated fatty acids.

The TG fatty acid carboxyl signals are dispersed according to their position in the glycerol moiety and their degree of saturation, as shown by Figure 1b and Supplementary Figure 1b. For the TG standard, which is composed of a mixture of synthetic triglycerides each with a single fatty acid species, the carboxyl signal profiles corresponding to SFA, MUFA and PUFA have identical aspects for both *sn*1,3 and *sn*2 positions as would be expected (Supplementary Figure 1). In contrast, human TG samples showed marked differences in relative SFA, MUFA and PUFA signal intensities between *sn*1,3 and *sn*2 positions (Figure 1b). The *sn*2 position was significantly more enriched with MUFA compared to the *sn*1,3 sites, while SFA abundance was significantly lower in the *sn*2 position compared to that of *sn*1,3 (Supplementary Figure 3).

**Figure 3.**
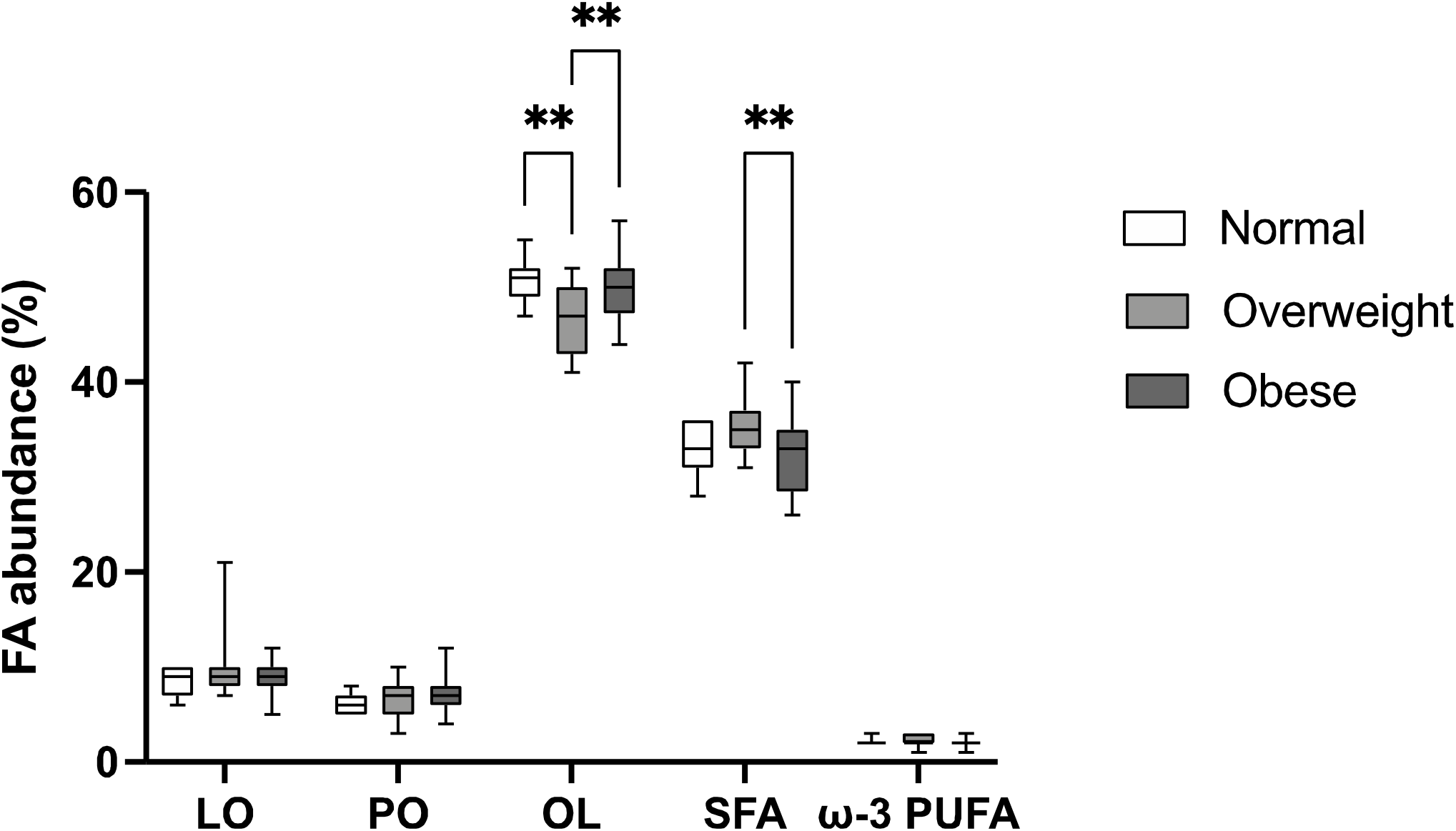
Triglyceride fatty acid composition of human adipose tissue biopsies based on analysis of the ω-3 ^13^C NMR signals and the fatty acid methyl ^1^H NMR signals. Samples categorized in weight status by BMI ranges. The fractional abundances of linoleate (LO), palmitoleate (PO), oleate (OL), saturated fatty acids (SFA) and ω-3 PUFA are shown. Normal n= 7, Overweight n= 19, Obese n=20. *p<0.05, ** p<0.006 (two-way ANOVA).

### Triglyceride fatty acid species and positional profiles in relation to BMI and HOMA-IR

^13^C NMR analysis of intact triglyceride informs the overall abundance of the different fatty acid constituents as well as their distribution between *sn*1,3 and *sn*2 glycerol positions (15,16). These characteristics were compared between subjects with normal, overweight and obese subjects categorized according to BMI and the results are shown in Figures 3 and 4. The OL fraction was significantly lower in overweight subjects compared to either normal or obese subjects while the SFA fraction was significantly higher in overweight compared to obese subjects (Figure 3). Regarding the distribution of different fatty acid classed in the TG glycerol moiety, the fraction of PUFA in the *sn*2 position was significantly lower in obese subjects compared to overweight subjects (p = 0.0472). The fraction of MUFA in the *sn*2 position was significantly higher in obese compared to overweight subjects (p = 0.0295), and also tended to be higher compared to the healthy subject group (Figure 4). Analysis of the fatty acid parameters as continuous variables for all subjects by linear regression analysis (Table 2, supplementary data) revealed significant positive correlations between the fraction of PO and BMI (R^2^ = 0.18, *p* = 0.0037) and the fraction of MUFA in the *sn*2 position and BMI (R^2^ = 0.12, *p* = 0.0164). There was also a significant negative correlation between the fraction of PUFA in the *sn*2 position and BMI (R^2^ = 0.11, *p* = −0.3339).

**Figure 4a,b.**
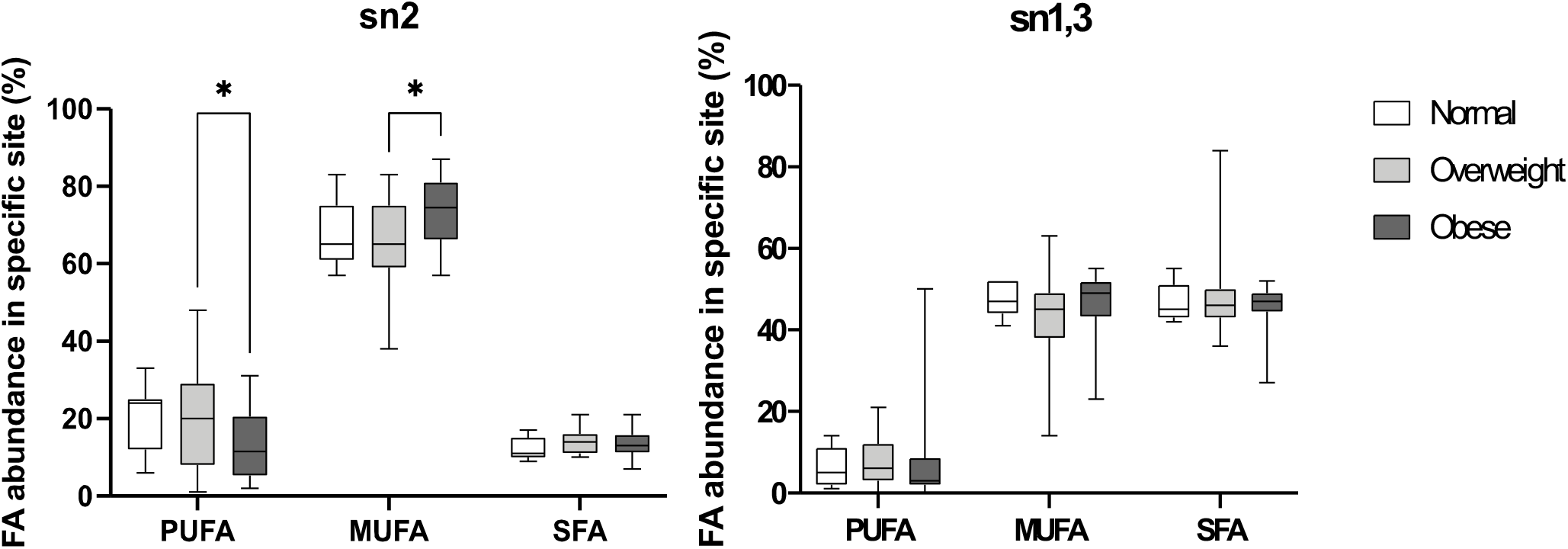
Distribution of SFA, MUFA and PUFA fatty acids in the *sn*1,3 (**a**) and *sn*2 positions (**b**) of the triglyceride in accordance with different BMI categories. Normal n= 7, Overweight n= 19, Obese n=20. *p<0.05 (two-way ANOVA).

As shown in Supplementary Table 3, the PO fraction was the sole parameter to be significantly correlated with HOMA-IR scores (R^2^ = 0.25, *p* = 0.0004).

## Discussion

Extensive studies have been performed to characterize the FA composition in TG obtained from both animal and plant sources (19,20). The most common approach involves transesterification of the TG to yield a mixture of FA-methyl esters, which are then analyzed by GC-MS (20,21). In comparison with ^13^C NMR, this approach provides better resolution of individual fatty acid species, in particular C14-C20 SFA. In addition, GC-MS has higher sensitivity allowing quantification of less abundant fatty acid species. However, it does not inform the distribution of fatty acids among the glycerol positions. More recently, LC-MS has been used to characterize the fatty acid composition of intact TG by measuring the molecular mass of the parent molecule and that of selected fragments following ionization (22). As for GC-MS, this methodology does not inform the positional distribution of different FA classes within the TG molecule. With ^13^C NMR, the main fatty acid classes accounting for 90-95% of the total TG fraction can be quantified, with resolution of some individual species, notably PO. There is currently high interest in PO since it has been found to be associated with obesity *per se* as well as with co-morbidities of obesity such as fatty liver and insulin resistance. In a study of overweight subjects, the molar fraction of PO in VLDL-TG was found to be positively correlated with elevated liver fat (23). In a cohort of nondiabetic subjects, Trico et al found that circulating non-esterified PO in the fasted state was positively correlated with fat mass but was also positively correlated with insulin sensitivity after adjustment for age, sex and adiposity (24). These and other studies support a possible role of PO as a beneficial lipokine: its release from adipose tissue improves insulin sensitivity in other tissues (25). However, this postulate is not supported by all studies. For example, Fabbrini et al. found no significant differences in the fraction of PO in neither VLDL nor NEFA between insulin-sensitive and insulin resistant subjects (26). In our study, we found that the levels of adipose tissue PO were positively associated with BMI but negatively correlated with insulin sensitivity. Gong et al. also reported a positive association between PO and BMI from subcutaneous adipose tissue collected from the upper buttock, (27). Interestingly, this relationship was strongest for individuals with the highest intake of sugar. To our knowledge, there have been no published studies that have examined the relationship between adipose tissue PO levels and whole-body insulin sensitivity. One explanation for adipose tissue and plasma PO to show opposite correlations with insulin sensitivity is that in the setting of insulin resistance, adipose tissue PO is somehow retained to a greater extent compared to other FA species during lipolysis thus generating a fasting non-esterified fatty acids (NEFA) profile that is relatively depleted in PO. Possible mechanisms could involve alterations in TG-FA cycling and/or changes in the selectivity of enzymes involved in TG synthesis and hydrolysis towards PO.

The positional distribution of FA in adipose tissue TG reflects the aggregated actions of the constituent enzymes involved in TG synthesis and turnover. During TG synthesis, FA are sequentially esterified at the *sn*1 and *sn*2 positions of glycerol-3-phosphate via glycerol-3-phosphate acyltransferase (GPAT) and acylglycerophosphate acyltransferase (AGPAT) to form phosphatidic acid (PA). Esterification at the *sn*3 site follows PA dephosphorylation to diacylglycerol and is catalyzed by diacylglycerol acyltransferase (DGAT). GPAT has a higher affinity for SFAs while AGPAT has a preference for mono- and polyunsaturated FAs over SFAs (10, 28). DGAT exists as two isoforms: DGAT1 and DGAT2. DGAT2 is postulated as the principal enzyme for endogenous TG synthesis while DGAT1 is involved in the resynthesis of TG from DAG during lipolysis (29) and may play a protective role in scavenging FA (30). The selectivity of either adipose tissue DGAT enzyme to different FA species is poorly characterized.

Lipolysis of TG involves sequential cleavage of FA by adipose triglyceride lipase (ATGL) to form diacylglycerol followed by hormone-sensitive lipase (HSL) and monoacylglycerol lipase (MGL). ATGL selectively hydrolyzes TG at the *sn*-2 position, with a high selectivity for PO. In the presence of the co-activator CGI-58, hydrolysis at the *sn*-1 position is also initiated resulting in the formation of *sn*1,3 and *sn*2,3 diacylglycerols, respectively (12). While HSL targets the *sn*3 position of diacylglycerol (31), it does not show particularly strong selectivity for different FA but may have some preference for unsaturated over saturated species (32). Given these various degrees of FA selectivity during TG synthesis as well as during cycling between TG DAG and MAG, it is not surprising that different FA species have non-uniform distributions in the glycerol positions.

Our data revealed that alterations in the distribution of MUFA and PUFA between the *sn*1,3 and *sn*2 sites of glycerol were associated with BMI independently of their total abundance. These data are consistent with differences in FA selectivity by one or more enzymes involved in TG metabolism and/or different TG-FA cycling activities in the setting of obesity. As previously discussed, such alterations could contribute to the adipocyte retention of PO in relative to other FA species. A key first step in testing this hypothesis would be to determine the relationship between adipose tissue and plasma NEFA PO composition in settings of obesity and insulin resistance.

In conclusion, we demonstrated that ^13^C NMR spectroscopy can be applied to analyze TG from adipose tissue biopsies of 12-65 mg. In addition to quantifying FA classes and selected species, ^13^C NMR also informs the positional distribution of FA within the TG molecule. Our study revealed for the first time a highly non-homogenous distribution of FA between the *sn*1,3 and *sn*2 sites of glycerol in human TG. Moreover, distributions of certain FA classes can be correlated with a particular phenotype or pathology – in this case obesity - independently of their overall abundance. This may reflect alterations in adipocyte TG-FA cycling and/or selectivity of enzymes involved in TG synthesis and hydrolysis.

## Abbreviations

(AGPAT): Acylglycerophosphate acyltransferase
(AT): Adipose tissue
(ATGL): Adipose triglyceride lipase
(BMI): Body mass index
(DGAT): Diacylglycerol acyltransferase
(FA): Fatty acids
(GPAT): Glycerol-3-phosphate acyltransferase
(HOMA IR): Homeostatic model assessment of insulin resistance index
(HSL): Hormone-sensitive lipase
(LO): Linoleate
(MTBE): Methyl-*tert*-butyl ester
(MGL): Monoacylglycerol lipase
(MUFA): Monounsaturated fatty acids
(NEFA): Non-esterified fatty acids
(OL): oleate
(PO): palmitoleate
(PUFA): Polyunsaturated fatty acids
(SFA): Saturated fatty acids
(TG): Triglycerides

