## Supplementary Table 1, Supplementary Table 2, Supplementary Table 3, Supplementary Figure 1, Supplementary Figure 2, Supplementary Figure 3 for "Positional and Compositional Analysis of Saturated, Monounsaturated and Polyunsaturated Fatty acids in Human Adipose Tissue Triglyceride by ^13^C NMR"

**Supplementary Material**

**Figure legends**

**Supplementary Figure 1.**  ^13^C NMR spectrum of a mixture of synthetic triglycerides featuring the ω, ω-2 and ω-3 signals for saturated fatty acids (SFA), oleic acid (OL) palmitoleic acid (PO) and linoleic acid (LO). SFA consisted of a mixture of myristate, palmitate and stearate.

**Supplementary Figure 2.**  ^1^H NMR spectrum of the mixture of synthetic triglycerides. The fatty acid methyl hydrogens are shown in expanded from and feature the ω-3 PUFA represented by α-linolenic acid (ω-3 CH_3_), the natural abundance ^13^C satellite signals of the other fatty acid methyl hydrogens (^13^C CH_3_) and the fatty acid methyl hydrogens bound to ^12^C (^12^C CH_3_).

**Supplementary Figure 3.**  Distribution of SFA, MUFA and PUFA in the *sn*1,3 and *sn*2 triglyceride positions of entire subject cohort. n=46 ***** p<0.001 (two-way ANOVA).*

**Supplementary Table 1.** ^13^C NMR chemical shift values for individual fatty acid species obtained from spectra of synthetic triglyceride standards. All chemical shifts are referenced to the center signal of the CDCl_3_ triplet which was assigned a value of 77.23 ppm.

|  | ***SFA*** | | | ***MUFA*** | | ***PUFA*** | | |
| --- | --- | --- | --- | --- | --- | --- | --- | --- |
|  | ***Myristate*** | ***Palmitate*** | ***Stearate*** | ***Oleate*** | ***Palmitoleate*** | ***Linoleate*** | | ***a-Linoleate*** |
| *𝜔 -1* | 14.34 | 14.34 | 14.34 | 14.33 | 14.33 | 14.29 | | 14.29 |
| *ω -2* | 22.92 | 22.92 | 22.92 | 22.90 | 22.86 | 22.80 | | 22.80 |
| *ω -3* | 32.15 | 32.15 | 32.15 | 32.13 | 32.01 | 31.75 | | - |
| *C1- sn2* | 173.10 | 173.10 | 173.10 | 173.06 | 173.06 | | 173.05 | 173.05 |
| *C1- sn1,3* | 173.51 | 173.51 | 173.51 | 173.48 | 173.48 | | 173.46 | 173.46 |

**Supplementary Table 2.** Simple linear regression analysis of triglyceride fatty acid fractions including those in the *sn*1,3 and *sn*2 positions from human AT biopsies *versus* BMI. A bivariate analysis (Pearson r) was applied to correlations that showed statistical significance *(*p* < 0.05).

|  | ***LO*** | ***PO**** | ***OL*** | ***SFA*** |
| --- | --- | --- | --- | --- |
| *R squared* | 0.02098 | 0.1763 | 0.02154 | 0.05683 |
| *P value* | 0.3368 | 0.0037 | 0.3304 | 0.1106 |
| *Pearson r* |  | 0.4199 |  |  |
|  | ***PUFA%- sn2**** | ***MUFA%- sn2**** | ***SFA%- sn2*** |  |
| *R squared* | 0.1115 | 0.1241 | 0.0002497 |  |
| *P value* | 0.0233 | 0.0164 | 0.917 |  |
| *Pearson r* | -0.3339 | 0.3523 |  |  |
|  | ***PUFA%- sn1,3*** | ***MUFA%- sn1,3*** | ***SFA%- sn1,3*** |  |
| *R squared* | 0.001611 | 0.01599 | 0.00955 |  |
| *P value* | 0.7911 | 0.4023 | 0.5182 |  |

**Supplementary Table 3.** Simple linear regression analysis of triglyceride fatty acid fractions including those in the *sn*1,3 and *sn*2 positions from human AT biopsies *versus* HOMA IR. A bivariate analysis (Pearson r) was applied to correlations that showed statistical significance *(*p* < 0.05).

|  | ***LO*** | ***PO**** | ***OL*** | ***SFA*** |
| --- | --- | --- | --- | --- |
| *R squared* | 0.004555 | 0.2492 | 0.004183 | 0.008213 |
| *P value* | 0.6558 | 0.0004 | 0.6694 | 0.5492 |
| *Pearson r* |  | 0.4992 |  |  |
|  | ***PUFA%- sn2*** | ***MUFA%- sn2*** | ***SFA%- sn2*** |  |
| *R squared* | 0.01589 | 0.0109 | 0.004588 |  |
| *P value* | 0.4039 | 0.4899 | 0.6547 |  |
|  | ***PUFA%- sn1,3*** | ***MUFA%- sn1,3*** | ***SFA%- sn1,3*** |  |
| *R squared* | 0.005458 | 0.01456 | 0.002754 |  |
| *P value* | 0.6256 | 0.4244 | 0.7291 |  |

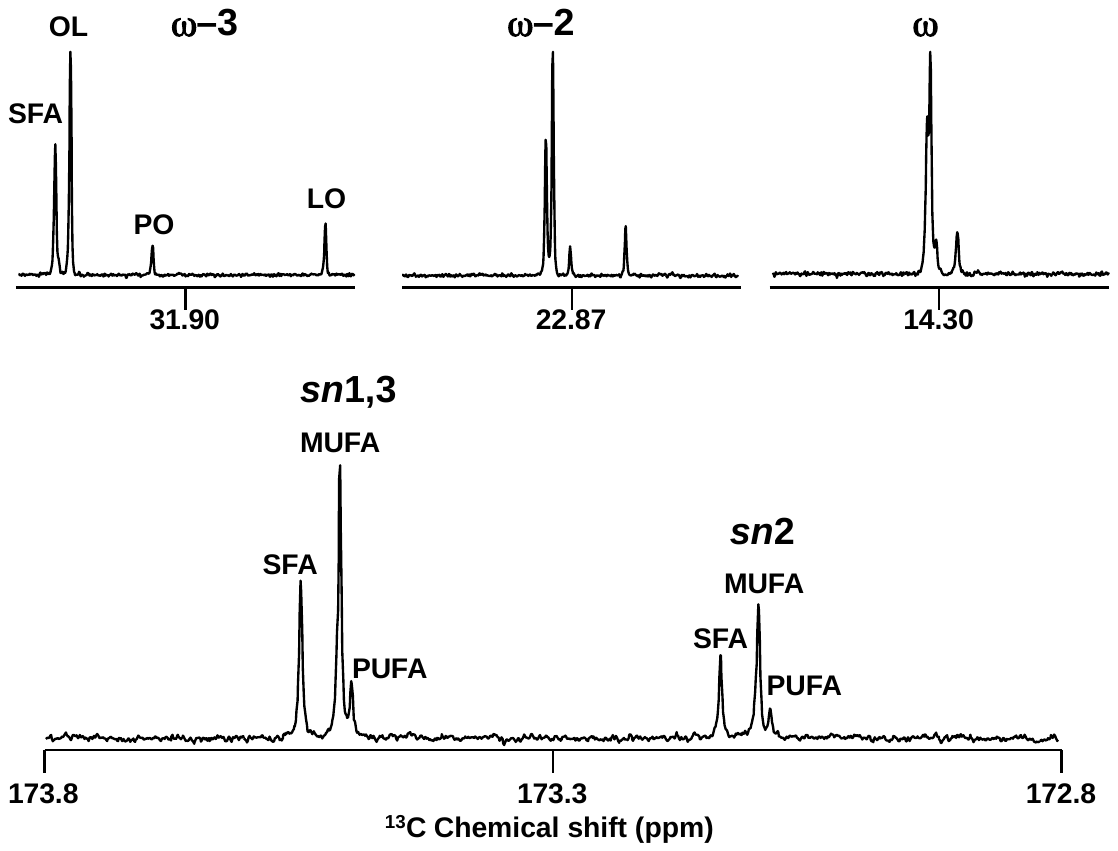

Supplementary Figure 1

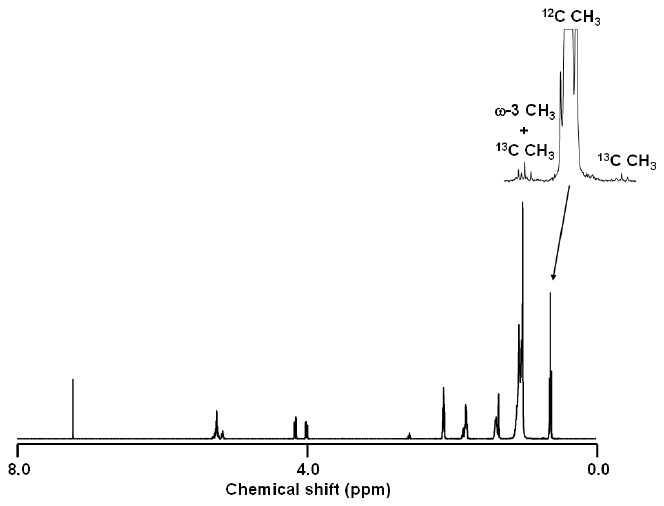

Supplementary Figure 2

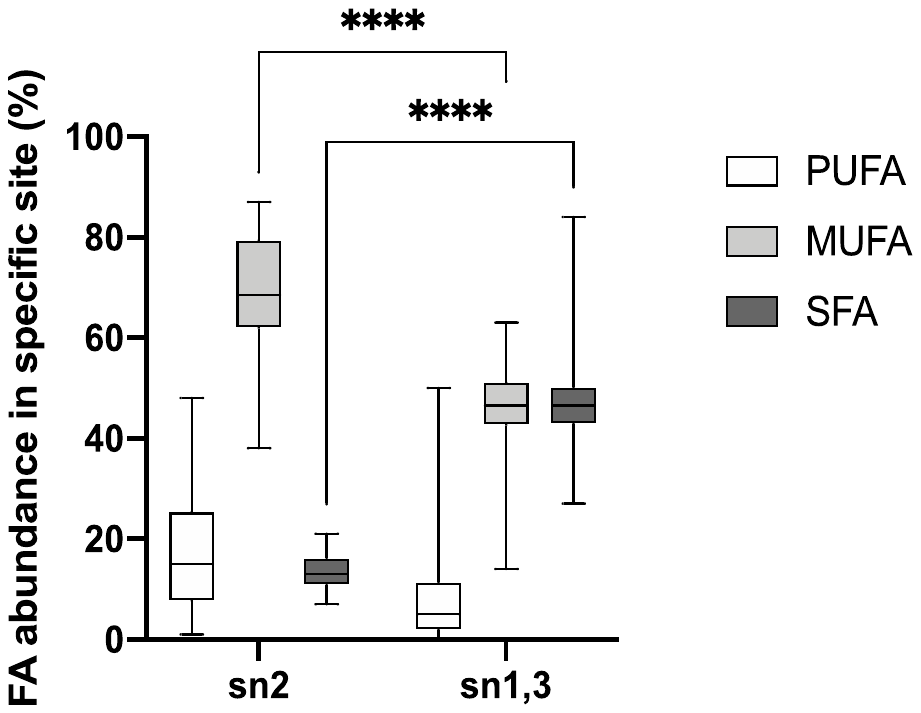

Supplementary Figure 3
